# Altered *glr-1* expression is linked to EtOH cessation-mediated behavior deficits in *C. elegans*

**DOI:** 10.1101/2025.11.19.689356

**Authors:** Katie Brandel-Ankrapp, Rachel N. Arey

## Abstract

Chronic ethanol (EtOH) exposure and withdrawal are linked to worsened memory and cognitive outcomes. One target of EtOH is the glutamatergic α-amino-3-hydroxy-5methyl-4-isoxazoleproprionic acid receptor (AMPAR). Both EtOH exposure and cessation influences AMPAR expression, function, and broad glutamate signaling. Studies of EtOHrelated AMPAR regulation suggest that EtOH regulation of AMPARs drives behaviors related to dependence and memory phenotypes. However, due to the complexity of the mammalian brain, it is difficult to unravel the precise mechanism by which EtOH regulates AMPARs in mammals to modify specific memory behaviors. In *C. elegans*, GLR-1, an AMPAR ortholog, has tightly defined expression in relatively few neurons, has conserved regulatory mechanisms, and is linked to molecularly conserved associative behaviors including those disrupted by EtOH. Using an established paradigm that exposes worms to chronic EtOH and cessation (“withdrawal”), we examined the relationship between cessation from chronic EtOH exposure, associative memory, and GLR-1/AMPAR regulation. We found that cessation from chronic EtOH disrupts intermediate-term associative memory (ITM) in wild-type worms, phenocopying loss of GLR-1 function. Next, we found that loss of GLR-1 occludes the ITM deficit and EtOH-ceased wild-types phenocopy *glr-1* loss-of-function in other glutamate-dependent behaviors. We demonstrate that cessation downregulates *glr-1*, and increases CREB-mediated transcriptional activity in neurons, whereas losing CREB protects against cessationinduced *glr-1* downregulation and associated behavior deficits. Overall, CREB-mediated repression of *glr-1* may regulate behavior deficits in cessation.

**Significance statement:** Animals must navigate and remember their environments to survive. The neuronal processes underlying these behaviors can be disrupted by chronic EtOH. EtOH interacts with many processes and pathways in the complex mammalian nervous system. Here, we established a paradigm combining chronic EtOH exposure with associative memory conditioning in *C. elegans* to study the consequences of EtOH exposure and cessation on memory behaviors and their underlying molecular pathways. We found cessation leads to memory deficits that depend on functional excitatory transmission and master transcriptional regulator, CREB.

## INTRODUCTION

Chronic alcohol, specifically ethanol (EtOH) exposure and cessation are associated with persistent changes in the brain from the level of neuronal physiology (Hedayati Moghadam et al., 2024; Lovinger and Roberto., 2025; Abrahao et al., 2017; Chandler et al., 2003; Popova et al.,2023) to organismal behavior, including memory (Burnett et al., 2016; Rose et al., 2023; Ambrose et al., 2001; Larnerd et al., 2023; Lee et al., 2008; Lindsay et al., 2022; Davies et al., 2004; Lee et al., 2009; Mira et al., 2019). Chronic EtOH exposure is linked to worsened cognitive outcomes in mammals, (Farr et al., 2005; Hoffman et al., 2019; Nunes, Kipp, and Reitz, 2019; Kipp et al., 2021; Fernandez et al., 2017; Sun et al., 2023), including in humans (Sabia et al., 2018; Pitel et al., 2007a; Pitel et al., 2007b; Zinn et al., 2004; Noël et al., 2012; Sullivan and Pfefferbaum, 2019; Ambrose et al., 2001). Behavior and memory deficits are also observed during EtOH cessation (Läck et al., 2007; Stephens et al., 2005; Cannizzaro et al., 2019; Cheng et al., 2025, preprint; Qiao et al., 2018). To better understand this process, it is imperative to understand molecular changes underlying these memory deficits. One conserved memory regulator is the ion-gated glutamate receptor family of α-Amino-3-hydroxy-5methyl-isoxazolepropionic acid receptor (AMPARs). Mammalian AMPARs regulate excitatory neurotransmission and drive lasting structural changes at synapses to promote plasticity and memory (Sanderson et al., 2008; Torquatto et al., 2019; Kessels and Malinow, 2009). Previous studies across multiple models demonstrated a link between cessation from EtOH and alterations in AMPA/glutamate signaling (Gerace et al.,2021; Christian et al., 2012; Wang et al., 2012; Wang et al., 2015; Gerace et al., 2019; Ulrichsen et al., 1996; Pickering et al., 2015; Das et al., 2016; Prisciandaro et al., 2019; Prisciandaro et al., 2016; Ende et al., 2013; Hermann et al., 2012; Mon et al., 2012; Thoma et al., 2011). However, the molecular mechanisms underlying altered glutamate receptor signaling during cessation, and their potential links to memory deficits remain unclear.

Barriers to addressing this question in mammalian models include heterogeneity of AMPA-receptor expressing neuronal subtypes within memory regulating circuits, and the labor-intensive nature of manipulating specific AMPA-regulating pathways in the context of ethanol exposure, cessation, and behavioral assays. Indeed, a reductionist model with a simple nervous system, conservation of responses to ethanol (both molecular and behavioral) and associative behaviors, would be advantageous.

Recently, the nematode *C. elegans* has been a useful system to untangle the molecular substrates of behavioral responses to ethanol. *C. elegans* have a stereotyped nervous system consisting of only 302 neurons (White et al., 1986; Witvliet et al., 2021). This simple nervous system coordinates conserved behavioral responses to relevant concentrations of ethanol including sensitivity, tolerance, and cessation/cessation. The genes underlying ethanol responses are conserved between worms and mammals, including humans (Mitchell et al., 2010; Thorsell 2007; Kapfhamer et al., 2008; Bettinger and Davies, 2014; Mathies et al., 2015; Davies et al., 2003; Davies et al., 2004).

In addition to EtOH-related behaviors, worms display molecularly conserved forms of associative memory to various stimuli, including odors (Kauffman et al., 2010; Stein and Murphy 2014; Rose et al., 2003; Gharat et al., 2024; Morrison and van der Kooy, 2001; Kano et al., 2008; Stetak et al., 2009). For example, olfactory learning and memory performance require proper function of GLR-1 (Morrison and van der Kooy, 2001; Stetak et al., 2009; Vukojevic et al., 2012; Gharat et al., 2024), the closest related orthologue to mammalian AMPA-type subunits (Brockie et al., 2001; Hart et al., 1995; Maricq et al., 1995). Unlike in higher organisms, where glutamate receptors are widely dispersed (Shen et al., 2021), *C. elegans* express GLR-1 in relatively few neuron types (Maricq et al., 1995) that are linked to specific behavioral metrics, making the worm an attractive model to study the interplay between mechanisms of ethanol and cessation-related changes in AMPA regulation. In fact, EtOH exposure during non-associative memory training modulates GLR-1 and is associated with memory deficits in worms (Rose et al., 2023). However, it is unknown whether cessation from a chronic exposure would affect associative behaviors in the worm via GLR-1 regulation.

Here, we examined the relationship between cessation from chronic EtOH exposure, associative memory, and GLR-1/AMPAR regulation. We found that cessation from chronic EtOH disrupts intermediate-term associative memory (ITM) in wild-type worms, phenocopying loss of GLR-1 function. Next, we found loss of GLR-1 occludes this ITM deficit and, after EtOH cessation, wild-types phenocopy *glr-1* loss-of-function in other glutamate-dependent behaviors. We demonstrate that cessation downregulates *glr-1* and a molecular pathway implicating the involvement of CREB orthologue CRH-1. Finally, we report evidence that EtOH cessation enhances CREB-mediated transcriptional activity.

## METHODS

### *C. elegans* strains and maintenance

All strains were maintained at 20°C on 10cm plates made from standard nematode growth medium (NGM: 3g/L NaCl, 2.5g/L of Bacto-peptone, 17g/L Bacto-agar in milliQ water) or high nematode growth medium (HGM: 3g/L NaCl, 20g/L of Bacto-peptone, 30g/L Bactoagar in milliQ water). After autoclaving and allowing molten agar to cool slightly, we added 1mL/L cholesterol (5mg/mL in ethanol), 1mL/L 1M CaCl_2_, 1mL/L 1M MgSO_4_, and 25mL/L 1M potassium phosphate buffer (pH 6.0) (Brenner, 1974). Experiments were performed using NGM plates seeded with OP50 *E. coli* as the food source for ad libitum feeding.

Hypochlorite population synchronization was performed by collecting eggs from gravid hermaphrodites via exposure to an alkaline-bleach solution (85mL water, 15mL sodium hypochlorite, 5mL 5M NaOH), followed by repeated washing of collected eggs in 1mL of M9 buffer (6g/L Na_2_HPO_4_, 3g/L KH_2_PO_4_, 5g/L NaCl and 1mL/L 1M MgSO_4_ in milliQ water.

### Strains

Wild-types (N2 Bristol) were obtained from the Caenorhabditis Genetics Center (University of Minnesota, Minneapolis, MN).

Transgenic strains: CQ161 (*pCRE::GFP*) was first described in (Kimura et al., 2002) and available in the Arey lab.

Mutants: KP4 (*glr-1*(n2461) III); CZ3761 [*ptp-3(mu256*) II], YT17 [*crh-1(tz2)* III], CB1393 [*daf-8*(e1393) I] were obtained from the Caenorhabditis Genetics Center (University of Minnesota, Minneapolis, MN). RNA58 (*glr-1*;*crh-1*) mutants were generated by KB-A by crossing KP4 [*glr-1(n2461)*] with YT17 [*crh-1(tz2*)].

### EtOH Treatment

Ethanol treatment of worms was performed as previously described (Lee et al., 2009;Salim et al., 2022), in which EtOH was added to 10cm NGM growth plates by pipetting EtOH onto the agar, wrapped in parafilm to prevent evaporation, and incubated at 20C for 2h . Worms were pipetted onto each plate and left to incubate at 20C for 2024h. For the cessation group, worms were washed from EtOH plates after incubation and transferred onto normal NGM plates for 1h, which has been published as sufficient time for internal EtOH concentrations to reach negligible levels (Scott et al., 2017).

### EtOH Preference Assay

EtOH preference assays were performed as previously described (Lee et al., 2009). Following exposure to water, EtOH, or EtOH cessation, 100-200 worms were washed with M9 and pipetted in the center of an unseeded 100mM plate that is separated into quadrants. Quadrants on opposite ends of the plate had either EtOH or water pipetted into a 9mm-sized well of the plate to reach a final concentration of 300mM. Plates were sealed with parafilm and EtOH was allowed to equilibrate in the agar for 2h prior to the start of the assay. Following 30 minutes after pipetting worms onto assay plates, worms were counted in each quadrant of the plate. The EtOH preference index was calculated as: {(#WormsEtOH quadrants – (#Wormscontrol quadrants)/#totalworms}

### Baseline chemosensation assays

Chemotaxis, or chemosensation experiments, were performed based on previously published assays (Bargmann et al., 1993). In brief, assays were performed on unseeded 10cm NGMs. Two marks were made on the back of the plate on opposite sides of the plate, approximately 0.5cm from the edge. 1μL of sodium azide (Thermo Fisher) was placed on both spots and allowed to dry before adding 1μL of test odorant diluted in ethanol or water on one side and ethanol on the other. The test odorant used was 10% butanone (vol/vol, Sigma Aldrich). Worms were washed off their plates and subsequently washed three times with M9 buffer, then placed near the bottom center of the plate, equidistant between the two marks, and allowed to chemotax for an hour. Chemotaxis indices for each timepoint were calculated as:

Chemotaxis index = (#worms_odorant_—#worms_ethanol_)/(total#worms).

### Positive olfactory associative memory assays

Water, EtOH-treated, or cessation-treated worms were trained and tested for intermediate-term memory changes as previously described (Kauffman et al., 2010; Kauffman et al., 2011; Hayden et al., 2024). Briefly, synchronized Day 2 adult worms were washed off plates with M9 buffer. Worms were then allowed to settle by gravity and washed twice more with M9 buffer to remove any bacteria. After washing, the worms were starved for one hour in M9 buffer. For 1x food-butanone pairing, hereby called conditioning, starved worms were transferred to 10cm NGM conditioning plates seeded with OP50 E. coli bacteria and with a total of 16μL of 10% butanone (Sigma Aldrich) diluted in ethanol streaked on the lid in a ‘#’ shape for one hour. After conditioning, the trained population of worms were tested for chemotaxis to 10% butanone (in ethanol or water) and to an ethanol control using standard, previously described chemotaxis conditions (Bargmann et al., 1993). Different stages of memory were tested by measuring chemotaxis of different subpopulations of worms at different timepoints at molecularly distinct stages of memory (Stein and Murphy, 2014). These stages are immediately after training (0min, learning) or after being transferred to 10cm NGM plates with fresh OP50 for 30 minutes (short-term associative memory), 1 hour (intermediate-term associative memory), or 2 hours (forgetting).

Chemotaxis indices for each timepoint were calculated as described earlier. Performance index is the change in the chemotaxis index after training relative to the untrained chemotaxis index, or:

**Performance index = (Chemotaxis index_trained)_—(Chemotaxis index_untrained_) Local search assays**

Following treatment of water, EtOH, or EtOH cessation, worms were picked onto unseeded NGM plates and allowed to acclimate for 5 minutes. Worms were video recorded for 5 minutes, keeping within the time frame of the local search period (Gray et al., 2005), and reversals and omega turns were manually scored. Recordings occurred over several days to ensure replication of results.

#### RNA Isolation, cDNA synthesis, and qRT-PCR

Worms of a particular genotype were crushed in liquid nitrogen and added to Trizol (Thermo Fisher Scientific). RNA was isolated per manufacturer’s instructions, followed by DNase treatment (Qiagen). cDNA was synthesized with an oligo dT primer and Superscript III reverse transcriptase enzyme (Thermo Fisher Scientific). cDNA was mixed with buffers, primers, SYBR green, and hot start Taq polymerase in a master mix prepared by a manufacturer (Thermo Fisher Scientific). Using a Quant Studio 7 Pro Dx Real-Time PCR System (Thermo Fisher Scientific), PCR reactions were run followed by a dissociation reaction to determine specificity of the amplified product. The amount of gene expression was quantified using the ΔΔCt method using pmp-3 as a reference gene.

Primer sets were as follows:

*pmp-3* For: 5’- AGTTCCGGTTGGATTGGTCC -3’

*pmp-3* Rev: 5’- CCAGCACGATAGAAGGCGAT-3

*glr-1* For: 5’-CTGTTGACGGGTCATCTGCT-3’

*glr-1* Rev: 5’-AACTGCACCTCCTTCGACTG-3’

### Confocal microscopy

Worms were paralyzed with fresh 4% levamisole diluted in M9 buffer and placed on a 4% agarose pad melted onto a microscope slide and secured with a coverslip. In all experiments, imaging of Day 2 adult worms was performed on a Nikon Ti2E inverted microscope system with a W1 spinning disk confocal unit at 40x. An excitation wavelength of 488nm for GFP with 50.8% laser power was used at 400ms exposure, with a pixel size of 0.18 µm/pixel, bit-depth of 16bit, and z-step size of 0.6 µm.

#### Image processing

The same settings were used for all conditions. All images were processed in FIJI to quantify fluorescence intensity and ROI detection (Schindelin et al., 2012). For segmentation of ROIs, z-stack images were transformed into maximum projections, set to 8-bit prior to thresholding with Otsu method, and watershed segmented to separate ROIs that were in proximity. The ROIs were collected in FIJI’s Particle Analyzer plugin and projected onto SUM projections from the original z-stacks for intensity quantification. Mean intensities collected from the SUM projection images were subtracted by the average mean intensity of three background areas within the worm. Because the GFP signal in negative controls is relatively dim to positive controls, representative images have enhanced brightness/contrast to see the outline of the worm.

### Statistical analysis

Statistical data is reported in the main text, figures, and tables as noted. Significance threshold of p < 0.05 was used. The symbols *, **, ***, and **** refer to p < 0.05, 0.01, 0.001, and 0.0001, respectively. For the comparison of performance indices between two behavior conditions (e.g. water-treated vs EtOH-treated), Welch’s t-test was used because it does not assume equal standard deviations. For comparison of performance indices between three or more groups (e.g. wild-type vs two different EtOH exposures), one-way analysis of variances followed by Bonferroni post hoc tests for multiple comparisons were performed. Two-way ANOVAs or mixed models were used for evaluating effects between condition (water-treated, EtOH, cessation) and timepoint (0hr, 0.5hr, 1hr, 2hr) on performance indices with a significant interaction between factors (p<0.0001) prompting Bonferroni post-hoc analyses to determine differences between individual groups. All experiments were repeated on separate days with separate populations to confirm reproducibility of results.

### Statistical analysis software

All statistics and code were run in GraphPad Prism 10, using standard toolboxes.

## RESULTS

### Cessation from chronic EtOH, but not chronic EtOH itself, disrupts intermediate term associative memory in wild-type worms

To assess the effect of EtOH cessation on associative learning and memory, we used a well-established positive olfactory association assay (Kauffman et al., 2010) in wild-type worms treated with 1 Water, 2) Chronic EtOH (20-24h) (**Figure 1B, left**) and 3) Chronic EtOH followed by 1h EtOH cessation, which we termed “Removal” in our figure legends (**Fig. 1B, right**). The memory assay trains worms to form a short-lasting positive association with a neutral odor (10% butanone) after pairing that odor with food for 1h (mass training) following a food deprivation period (**Figure 1A, left**). Learning and memory are quantified by measuring worm chemotaxis to butanone at different time-points after training (**Fig 1A, right**). Each learning and memory time-point is molecularly distinct and requires components needed for analogous memory stages across phyla (Kauffman et al., 2010; Stein and Murphy 2014). Previous studies tested chemotaxis and olfactory associative memory by providing two choices: 1) A control odorant, ethanol (weak attractant) and 2) A test odorant, typically diluted in ethanol. (Bargmann et al., 1993; Kauffman et al. ,2010; Kauffman et al., 2011; Hayden et al., 2024; Stein and Murphy 2014, Arey et al., 2018; Arey et al., 2019). Interestingly, exposing worms to high external concentrations of EtOH can yield EtOH preference (Lee et al., 2009). Based on this, we hypothesized that our EtOH exposure protocol could lead to EtOH attraction, which could introduce a confounding variable in our measure of a positive association formed with the odor diluted in ethanol. Prior to chemotaxis experiments, we exposed adult Day 1 (D1) worms to a high external concentration of EtOH (400mM) on their growth and maintenance plates for 20-24h. This external concentration yields an internal concentration of ∼50mM in worms after 24h (Scott et al., 2017), which equates to an intoxicating blood-EtOH concentration in humans (∼0.2%). Moreover, this concentration alters multiple behaviors and induces non-associative memory deficits when animals are exposed during training (Rose et al., 2023). In agreement with previous studies, we found worms tested immediately after 24h of exposure had significantly increased preference to EtOH vs. water-treated and EtOH-cessation-treated worms (**Extended Data Fig. 1-1A**).

**Figure 1:**
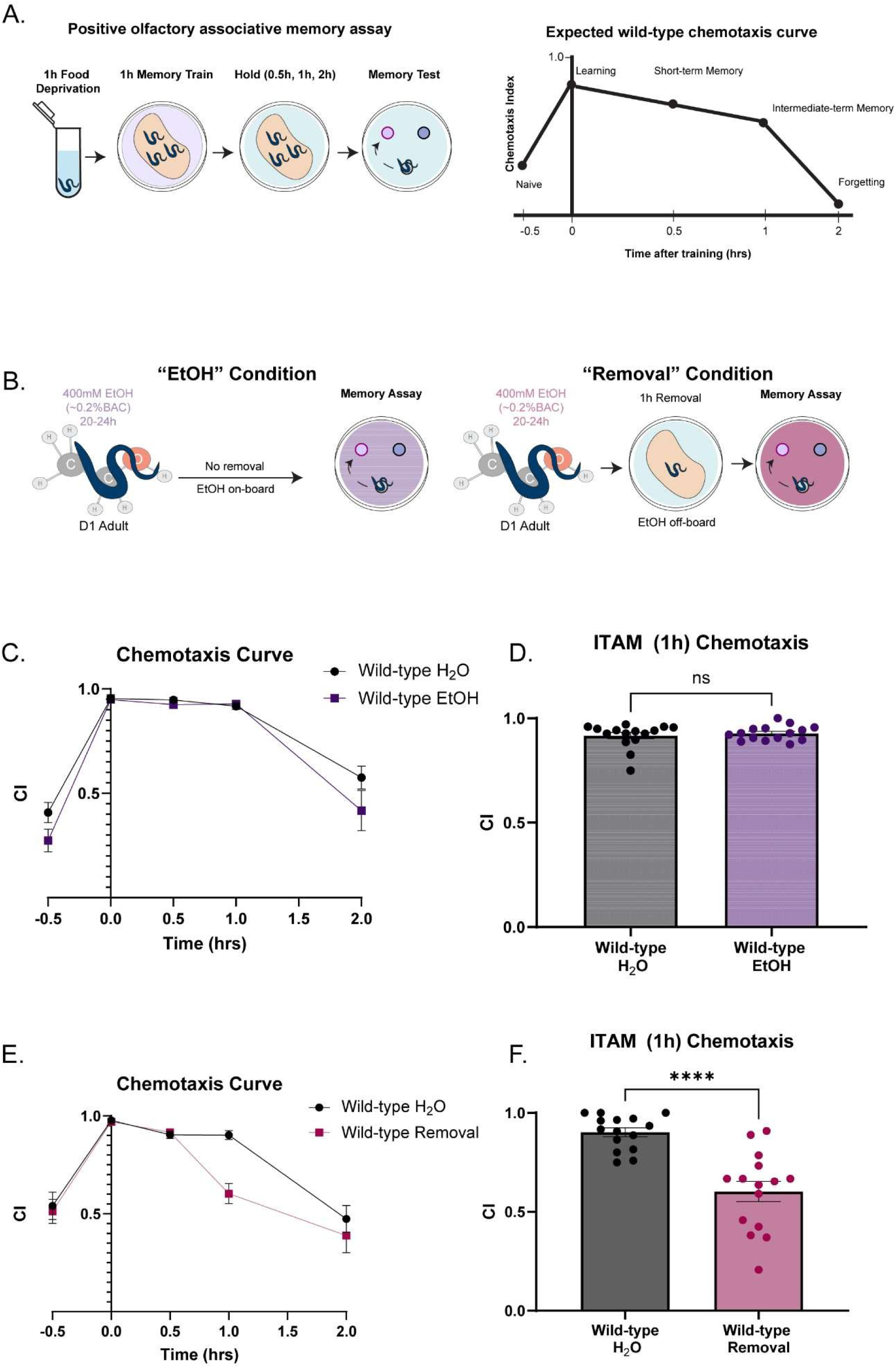
Cessation from chronic EtOH disrupts ITM. A) Schematic of experimental timeline for the positive olfactory associative memory assay. Briefly, young adult worms are food-withdrawn for 1h, then transferred onto food plates with 16uL of 10% butanone in water streaked onto the top of the plate. Worms incubate on these plates for 1h (associative memory training). Following 1h, worms were transferred onto food plates without the odor for 30 mins, 1h, and 2h prior to chemotaxis assays. For the T0 time-point, worms are immediately transferred from training plates onto chemotaxis assay plates. B) Expected chemotaxis curves of wild-type worms for every time-point in the assay. “On board” is defined as EtOH still being present in the worms by the start of the assay, as they did not undergo the 1h cessation/removal period. “Off board” is defined as EtOH no longer being present in the worms by the start of the assay, due to undergoing the 1h cessation/removal period. C) EtOH exposure schematic for each condition. Young-adult worms were transferred onto food and EtOH-treated plates for 20-24h. The “EtOH” condition immediately continues to the experiment, while the cessation condition is transferred onto normal food plates for 1h prior to experiments. To control for washes and transfers experienced by worms upon EtOH cessation, water- and EtOH-treated worms are also washed and transferred onto their appropriate plates. D) Chemotaxis curve of EtOH-treated and water-treated wild-types across all time points the associative memory assay (Mixed-effects analysis: Time x Condition, F(1.842, 51.10)=1.684, p=0.1648; Bonferroni’s multiple comparisons test: Naïve:Mean_Water_=.4077, Mean_EtOH_=.2749 (95% CI: -.06879,.3366),t(26.41)=1.833,p=.3902;0h: Mean_Water_=.9535,Mean_EtOH_=.9484(95% CI:-.05499,.06519),t(20.67)=.2406, p>0.9999; 0.5h: Mean_Water_=.9472, Mean_EtOH_=.9281, (95% CI:-.02807,.07251), t(26.91)=1.224, p>0.9999; 1h:Mean_Water_=..9171, Mean_EtOH_=.9281 (95% CI: -.06080,.03885),t(22.83)=.6189, p>0.9999; 2h:Mean_Water_=.4169 Mean_EtOH_=.1583, (95% CI: -.1542,.4707),t(22.21)=1.426, p=0.8384). N=14-15 plates per condition. N.s., not significant (p>.05) Error bars are SEM. E) Chemotaxis index of the 1h time-point (Welch’s t-test:Mean_Water_=.9171, Mean_EtOH_=.9281, (95% CI: -.02573-.04768), t(22.83)=.6189, p=0.5421). N=15 plates per condition. N.s., not significant (p>.05). F) Chemotaxis curve of and water-treated and cessation-treated wild-type worms across all time points in the associative memory assay (Mixed-effects analysis, Time x Condition, F(4,111)=3.358, p=0.0123; Bonferroni’s multiple comparisons test: Naïve:Mean_Water_=.5403, Mean_Cessation_=.5132(95% CI:-.2318,.2861, t(26.84)=.2907, p>0.9999; 0h:Mean_Water_=.9772, Mean_Cessation_=.9705(95% CI:-.03700,.05042),t(26.09)=.4264, p>0.9999; 0.5h:Mean_Water_=.9040, Mean_Cessation_=.9159 (95%CI:-.08483,.06098), t(27.98)=.4521,p>0.9999; 1h:Mean_Cessation_=.9021,MeanWildtype+Cessation=.6026,(95% CI:.1392,.4598), t(19.13)=5.340,p=0.0002;2h: Mean_Water_=.4738, Mean_Cessation_=.3885,(95% CI: - .2233,.3939),t(26.48)=.7670, p>0.9999. N=14-15 plates per condition. N.s., not significant (p>.05). Error bars are SEM. F) Chemotaxis index of the T60 time-point (Welch’s t-test: Mean_Water_=.9021, Mean_Cessation_=.6026, (95% CI: -.4168,.1822), t(19.13)=5.340, p<.0001). N=15 plates per condition. ****p<.0005. Error bars are SEM.

We next assessed if we could instead dilute butanone in water for our behavioral assays. We found that wild-type animals exhibit normal chemotaxis when butanone is diluted in water (**Extended Data Fig. 1-1B**); therefore, all subsequent behavior experiments were performed with water as the diluent instead of EtOH.

With conditions for assessing behavior in the context of ethanol exposure optimized, we next compared learning and memory performance between young-adult wild-type worms following each treatment. To withdraw worms from EtOH, worms are washed with M9 and transferred onto normal plates for 1h, as internal concentrations drop to negligible levels at this time-point (Scott et al., 2017). Neither chronic EtOH nor EtOH cessation significantly altered baseline chemosensation (**Extended data Fig 1-1C, D**). Worms that were treated with chronic EtOH had comparable chemotaxis indices to water-treated controls across all time-points in the assay, including the ITM time-point (**Fig. 1C, D**). This suggests that chronic EtOH treatment itself does not lead to deficits in positive associative learning and memory. However, worms that underwent the 1h cessation period prior to the assay performed worse than water-treated controls specifically at the intermediate term memory time-point while learning and short-term memory were spared (**Fig. 1E, F**). Overall, we discovered an EtOH-cessation-dependent deficit in positive olfactory associative intermediate-term memory consolidation.

### Cessation from chronic EtOH interacts with an AMPAR orthologue, GLR-1

Previous studies demonstrate that learning and memory in *C. elegans* requires GLR-1 (Gharat et al., 2024; Rose et al., 2003; Morrison and van der Kooy, 2001; Kano et al., 2008; Stetak et al., 2009; Vukojevic et al., 2012; Hadziselimovic et al., 2014) which led us to hypothesize that cessation from chronic EtOH interacts with GLR-1 to lead to ITM deficits. We confirmed that loss of GLR-1 results in memory deficits in our assay. *glr-1* mutant animals exhibit normal chemotaxis, and learning, but have significantly reduced ITM when compared to wild-type animals (**Figs. 2A, B**). We next sought to examine the effects of EtOH cessation on *glr-1* mutant learning and memory. Because we were interested in studying the interaction between ethanol and GLR-1, we assessed if more severe behavioral deficits than those present in *glr-1* mutants could be observed. We tested the behavior of *ptp-3* mutants, which lack functional PTP-3A, a receptor tyrosine phosphatase that regulates GLR-1 dynamics and perform poorly learning and memory (Pierce et al., 2022). We found *ptp-3* mutants have a subtle learning deficit **(Extended data Figs 2-1B**) but exhibit poorer ITM than both wild-types and *glr-1* mutants (**Fig.2A, B**). We then evaluated the behavior of *glr-1* animals after EtOH cessation. We found that EtOH-ceased *glr-1* mutants performed similarly to water-treated mutants (**Fig. 2C, D**) and did not exhibit a change in ITM performance indices between treatments. This contrasts with wild-type animals that exhibited a decrease in ITM performance indices after EtOH cessation (**Fig. 2E**), suggesting an interaction between EtOH cessation and *glr-1* (non-ITM time-points are shown in **Extended data Figs. 2-1 C, D**).

**Figure 2:**
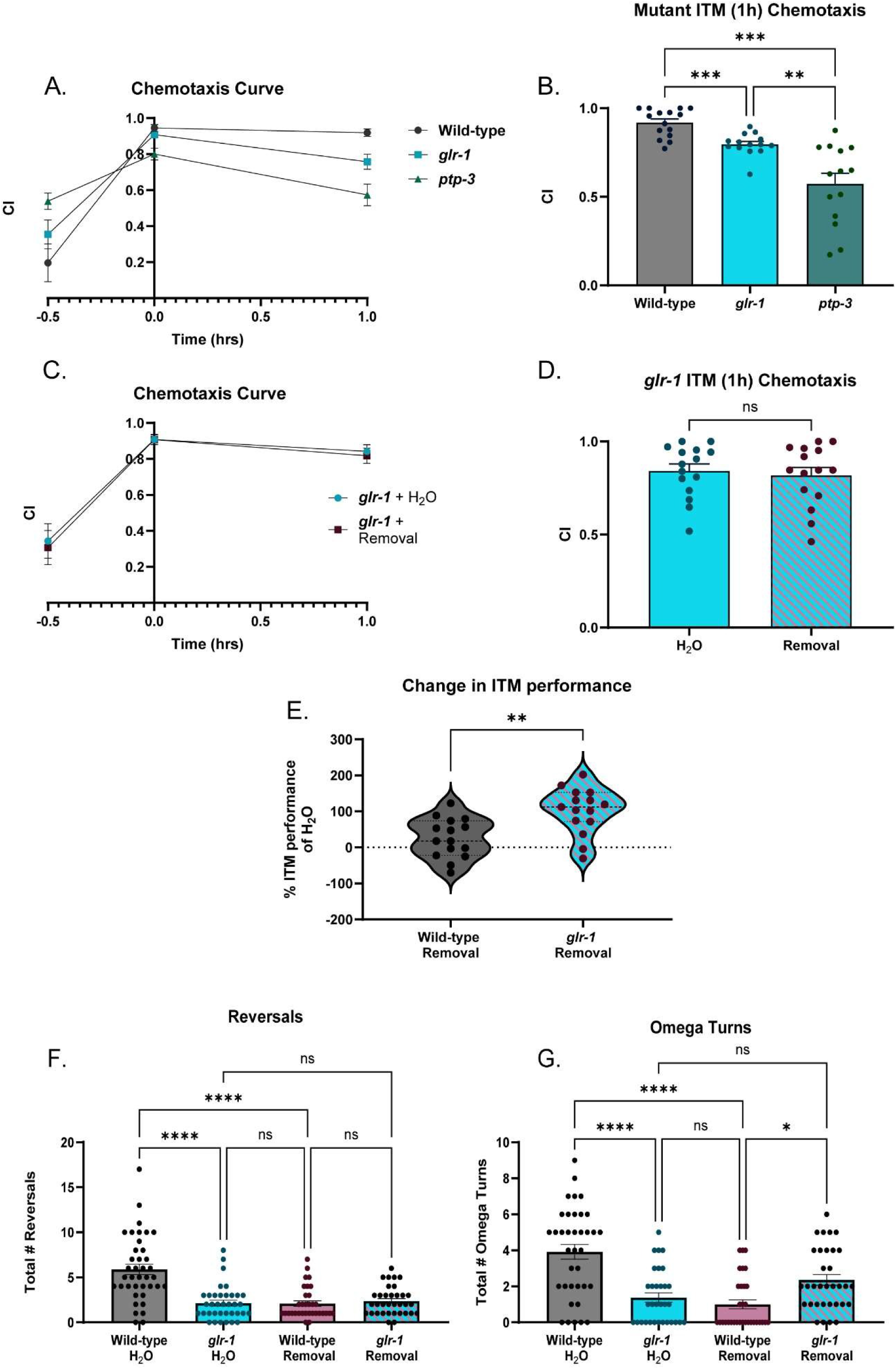
EtOH-cessation-treated wild-type worms phenocopy loss of *glr-1* function across multiple behaviors. A) Chemotaxis curves of wild-type vs. *glr-1* and *ptp-3* worms across time-points assessed in the associative memory assay. (Mixed-effects analysis, Time x Genotype: F(2.425,75.17)=10.45, p<.0001; Bonferroni’s multiple comparisons test, Naïve: Mean_Wild-type_=.1960 vs. Mean*_glr-1_*=.3548 (95% CI: .4964,.1790), t (26.13)=1.202, p=.72; Mean_Wild-type_=.1960 vs. Mean*_ptp-3_*=.5384 (95% CI:-.6425,-.4220), t(18.96)=2.995, p=.0224; Mean*_glr-1_*=.3548 vs. Mean*_ptp-3_*=.5384 (95% CI:-.4211,.05391), t(22.06)=2.003,p=.1730; 0h: Mean_Wild-type_=.9457 vs. Mean*_glr-1_*=.9082 (95% CI:.02849,.09989), t(25.11)=1.542,p=.4070; Mean_Wild-type_=.9457 vs. Mean*_ptp-3_*=.8006 (95% CI:.04585,.2445), t(21.66)=3.792),p=.0031; Mean*_glr-1_*=.9082 vs. Mean*_ptp-3_*=.8006 *glr-1* vs. *ptp-3*: (95% CI:.01362,.2017),t(17.64)=3.028,p=.0221; 1h: Mean_Wild-type_=.9189 vs. Mean*_glr-1_*=.7576 (95% CI:.03940,.2832), t(20.35) =3.452, p=.0074; Mean_Wild-type_=.9189 vs. Mean*_ptp-3_*=.5724 (95% CI:.1751,.5158), t(15.99)=5.421,p=.0002; Mean*_glr-1_*=.7576 vs. Mean*_ptp-3_*=.5724 (95% CI:-.005242,.3736),t(23.51)=2.506,p=.0586. N=15 plates per condition. N.s., not significant (p>.05), **p<.005,***p<.0005. Error bars are SEM. B) Chemotaxis index of wild-type vs. *glr-1* and *ptp-3* worms at ITM (1h) time-point: F=21.44(2.00, 18.31), p<.0001; (Brown-Forsythe ANOVA test, 21.44(2,18.31), p<.0001;Dunnett’s T3 multiple comparisons test: Wild-type vs. *glr-1*: Mean_Wild-type_=.9189, Mean*_glr-1_*=.7965, (95% CI: .05473,.1900), t(26.35)=4.603, p=.0003); Wild-type vs. *ptp-3*: Mean_Wild-type_=.9189, Mean*_ptp-3_*=.5734 (95% CI: .1766,.5143), t(15.99)=5.421,p=.0002), *glr-1* vs. *ptp-3*: Mean*_glr-1_*=.7965, Mean*_ptp-3_*=.5734 (95% CI: .05590,.3903), t(15.02)=3.561,p=.0083). N=14-15 plates per genotype. **p=.0083, ***p<.0005. Error bars are SEM. C) Chemotaxis curves of *glr-1* worms treated with water or EtOH cessation (Mixed-effects analysis: Time x Condition, p=0.8731, F(1.405,38.64)=.06824; Bonferroni’s multiple comparisons test: Naïve: Mean_Water_=.3436, Mean_Cessation_=.3068 (95% CI: -.3095,.3831), t(26.98)=.2713, p>.9999; 0h: Mean_Water_=.9077,MeanCessation=.9075 (95% CI: .1008,.1013), t(27.94)=.006104, p=.9952; 1h: Mean_Water_=.8423, Mean_Cessation_=.8180 (95% CI: -.12.01,.1687), t(27.37)=.4291, p=.6712). N=14-15 plates per condition. N.s., not significant (p>.05). D) Chemotaxis indices at ITM (1h) time-point in water or EtOH-cessation-treated *glr-1* worms. (Welch’s t test: Mean_Water_:.8423, Mean_Cessation_:.8180 (95% CI: -.1404,.09179) t(27.37)=.4291, p=.6712). N=14-15 plates per condition. N.s., not significant (p>.05). Error bars are SEM. E) Difference in performance in wild-type worms and *glr-1* worms treated with water or EtOH cessation (Welch’s t test, Mean_Wild-type_ =25.68, Mean*_glr-1_*=101.7 (95% CI: - 120.6,31.48), t(27.39)=3.499, p=.0016). N=15 plates per condition. **p<.005. Error bars are SEM. F) Reversals during local search behavior in water-treated wild-type and *glr-1* worms, and cessation-treated wild-type and *glr-1* worms (Kruskal-Wallis test: 38.40, p<.0001; Dunn’s multiple comparisons test, Mean rank Wild-type + Water =105.4, Mean rank *glr-1* + Water=56.50, Mean rank Wild-type + Cessation=55.48, Mean rank *glr-1* + Cessation=64.09. Wild-type + Water vs. *glr-1* + Water: Z=5.213, p<.0001; Wild-type + Water vs. Wild-type + Cessation: Z=5.200, p<.0001; Wild-type + Water vs. *glr-1* + Cessation: Z=4.338, p<.0001; *glr-1* + Water vs. Wild-type + Cessation: Z=.1039, p>.9999; *glr-1* + Water vs. *glr-1 +* Cessation: Z=.7825,p>.9999; Wild-type + Cessation vs. *glr-1* + Cessation: Z=.8682; p>.9999). N=33-39 worms per condition. N.s., not significant (p>.05), ***p<.0001. Error bars are SEM. G) Omega turns during local search behavior in water- or cessation-treated wild-types, *glr-1* worms(Kruskal-Wallis test: 34.81,p<.0001; Dunn’s multiple comparison’s test, Mean rank Wild-type + Water=93.91, Mean rank *glr-1* + Water=40.39, Mean rank Wild-type + Cessation=44.47, Mean rank *glr-1* + Cessation=73.79. Wild-type + Water vs. *glr-1* + Water: Z=4.400 p<.0001; Wild-type + Water vs. Wild-type + Cessation: Z=5.385, p<.0001; Wild-type + Water vs. *glr-1* + Cessation: Z=2.209, p=.1629; *glr-1* + Water vs. Wild-type + Cessation ,Z=.9516, p>.9999, *glr-1* + Water vs. *glr-1* + Cessation: Z=2.149, p=.1899; Wild-type + Cessation vs. *glr-1* + Cessation: Z=3.108, p=.0113). N=33-37 worms per condition. N.s., not significant (p>.05), *p<.05, ****p<.0001 Error bars are SEM.

Based on our findings that ethanol cessation appears to phenocopy *glr-1* reduction of function, and that *glr-1* mutants might exhibit resistance to EtOH cessation-related behaviors, we sought to further examine the potential relationship between EtOH cessation and GLR-1. To do this, we compared glutamate-dependent foraging behaviors between wild-type and *glr-1* worms across untreated control and 1h EtOH cessation conditions. These foraging behaviors, reversals and omega turns, are most prominent within minutes (1-12) of transferring worms from maintenance plates onto plates with no bacteria (Gray et al., 2005). Previous studies have shown loss of *glr-1* represses these behaviors (Juo et al., 2007; McGehee et al., 2015; Bhardwaj et al., 2020; Park et al., 2021; Hills et al., 2004; Luth et al., 2021; Campbell et al., 2016), whereas gain of *GLR-1* activity increases these behaviors (Zheng et al., 1999; Luth et al., 2021), providing a direct behavioral read-out of *glr-1* function. We found, as reported previously, untreated *glr-1* mutants performed fewer reversals and omega-turns than untreated wild-type animals.

Wild-type worms that received EtOH cessation also performed significantly fewer reversals than untreated wild-types, phenocopying *glr-1* worms, similar to what we observed with ITM (**Figs. 2F, G**).

### Cessation from chronic EtOH downregulates *glr-1*

Based on our combined behavioral data, we hypothesized that cessation from chronic EtOH led to decreased *glr-1* expression. We evaluated this at the transcript level using quantitative real-time PCR (qPCR) in wild-type worms receiving either water, chronic EtOH, or chronic EtOH + cessation. While *glr-1* transcript levels are unchanged relative to water controls in worms that were still on EtOH at the time of collection, 1h cessation resulted in a significant downregulation of *glr-1* transcript levels compared to other treatments (**Figs. 3A, B**). To assess if this downregulation was specific to *glr-1* glutamate receptors we also measured transcript levels of another type of ionotropic glutamate receptor, *nmr-1*, which is orthologous to mammalian N-methyl-D-aspartate (NMDA) receptor proteins. NMDA receptors are critical for plasticity and memory (Malenka and Nicoll, 1999) and are an EtOH target (Kumari and Ticku 2000; Morisot and Ron, 2017; Nagy, 2008; Wang et al.,2007b). Interestingly, we found that *nmr-1* transcript levels were not significantly different compared to water-treated wild-types in all exposure condition measured (**Fig. 3C**) These data suggest that a general downregulation of ionotropic glutamate receptors in nematodes does not occur after EtOH cessation, suggesting that EtOH cessation may specifically impact *glr-1* transcription, rather than widely repress ionotropic glutamate receptor transcription.

**Figure 3:**
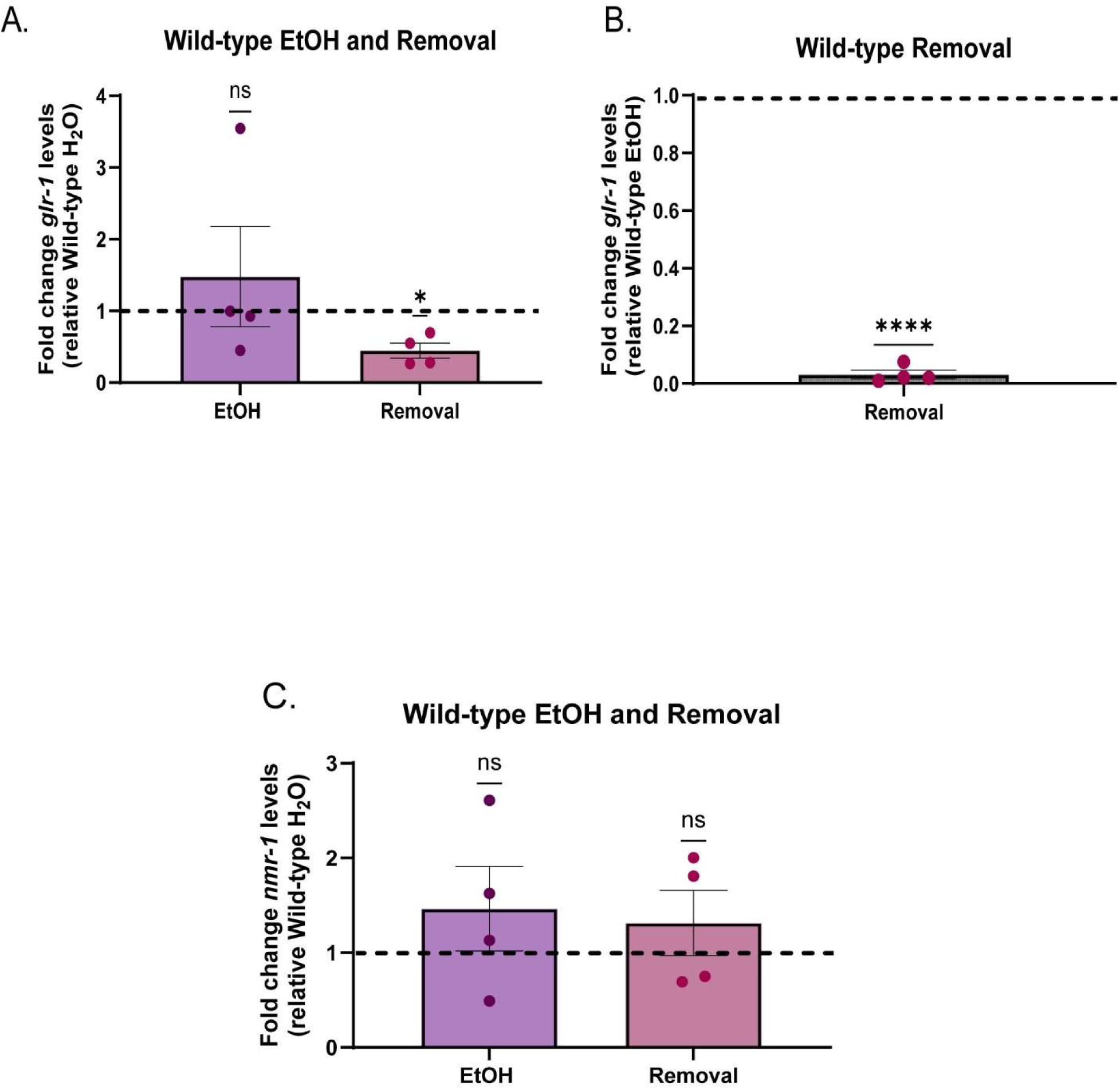
Cessation from chronic EtOH downregulates *glr-1*. A) Fold change in *glr-1* transcript levels between EtOH- and cessation-treated wild-types relative to water-treated wild-types (One sample t-test, EtOH: Theoretical mean=1, actual mean=1.480,(95% CI:.1.744,2.703), t(3)=0.6868,p=.5416; Cessation: Theoretical mean=1,actual mean=.4475, (95% CI:-.8869,-.2180), t(3)=5.257,p=.0134). N=4 samples per condition. N.s., not significant (p>.05), *p<.05. Error bars are SEM. B) Fold change in *glr-1* transcript levels in cessation-treated wild-types relative to EtOHtreated wild-types (One sample t-test, Theoretical mean=1.00, Mean=.03148, (95% CI: 1.016,-.9212), t(3)=65.15, p<.0001) N=4 samples per condition. ****p<.0001. Error bars are SEM. C) Fold change in *nmr-1* transcript levels between EtOH and cessation-treated wild-types relative to water-treated wild-types (One sample t-test, EtOH: Theoretical mean=1, actual mean=1.464 (95% CI:-.9558,1.884),t(3)=1.041,p=.3746; Cessation: Theoretical mean=1, actual mean=1.315,(95% CI:-.7819,1.411), t(3)=.9134, p=.4284 ). N=4 samples. N.s., not significant (p>.05). Error bars are SEM.

### Changes in *glr-1* expression in EtOH cessation may be CREB-dependent

We next sought to identify the molecular pathway underlying EtOH cessation-related glr1 downregulation we had observed. Previous studies have uncovered negative transcriptional regulators of *glr-1*, including those that are downstream of conserved molecular pathways in neurons such as DAF-7/TGF-β and CMK-1/CAMKIV (McGehee et al., 2015; Moss et al., 2016), which act through downstream transcription factor RSMAD/DAF-8 and CRH-1/CREB, respectively (**Fig. 4A**). Interestingly, alterations in both neuronal TGF-β and CREB expression have been documented in EtOH treatment in other models (Huang et al., 2015; McClinktick et al., 2020; Pandey et al., 2001; Bison and Crews 2003; Kurokawa et al., 2013; Shibasaki et al., 2011; Qiao et al., 2018; Wang et al., 2007a). Therefore, these pathways were attractive candidates for *glr-1* downregulation during EtOH cessation. To test this, we performed qPCR in either *daf-8* or *crh-1* mutant backgrounds following treatment with water or EtOH cessation. We hypothesized that if either *daf-8* or *crh-1* is involved in *glr-1* downregulation during EtOH cessation, then removing its ability to repress *glr-1* via genetic ablation would prevent glr1 downregulation in the context of cessation. In accordance with previous findings, we found that *daf-8* mutants had significantly higher *glr-1* transcript levels compared to wildtype worms (**Fig. 4B**) (McGehee et al., 2015). Following EtOH cessation, *daf-8* worms no longer had significantly higher *glr-1* levels relative to wild-types. Direct comparison of *daf-8* water-treated worms and *daf-8* EtOH-ceased worms was not significant (**Fig. 4B**), although the fold-change of *glr-1* transcript levels was significantly lower in *daf-8* worms after EtOH cessation relative to *daf-8* water-treated worms (**Figure 4C**). These results indicated that the downregulation in *glr-1* following EtOH cessation likely was not dependent on *daf-8*. When we assessed *glr-1* levels in *crh-1* mutants, we did not observe increased *glr-1* transcript levels as previously reported (Moss et al., 2016) (**Fig. 4D**). This discrepancy could be due to differences in life-stage, as we collected synchronized D2 adults for all experiments instead of mixed populations, or at the L4 stage that were used in previous work (Moss et al., 2016), or because we measured endogenous *glr-1* levels instead of a transcriptional *glr-1* reporter. However, we did find that *crh-1* worms did not exhibit a significant decrease in *glr-1* expression after EtOH cessation (**Fig. 4E**). These results are consistent with *crh-1* being a negative regulator of *glr-1* as previously reported in (Moss et al., 2016), albeit in a context-dependent manner in adulthood. Overall, these results suggest that a specific molecular pathway involving *crh-1* and CREB-mediated transcription is required for *glr-1* downregulation during EtOH cessation in *C. elegans*.

**Figure 4:**
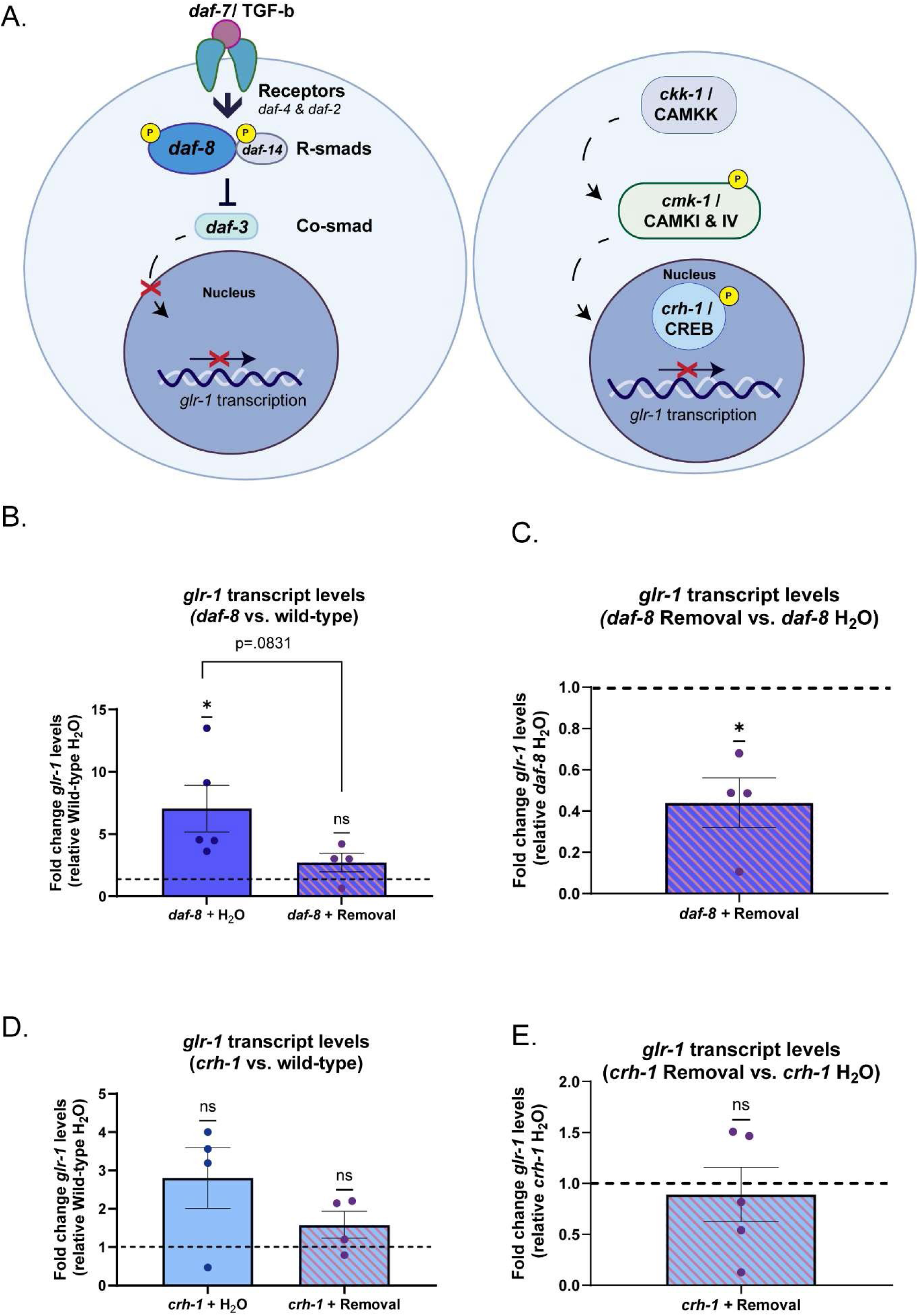
Changes in *glr-1* expression in EtOH cessation may be CREB-dependent. A) Schematic of candidate signaling pathways known to transcriptionally regulate *glr-1* downstream of DAF-7/TGF-β(left), and CMK-1/CAMK1 (right). B) Relative fold change of *glr-1* mRNA levels in *daf-8* background relative to wild-type controls. (One-sample t-test, *daf-8* + Water vs. Wild-type + Water: Theoretical mean=1, actual mean=7.054 (95% CI: .8324,11.27), t(4)=3.219, p=.0323; *daf-8* + Cessation vs. Wild-type + Water : Theoretical mean=1, actual mean=2.721 (95% CI:,-.6451,4.087), t(3)=2.315, p=.1036). Welch’s t test *daf-8* + Water vs. *daf-8* + Cessation. Mean*_daf-8_*_+Water_ = 7.054 Mean*_daf-8_*_+Cessation_= 2.721 (95% CI: -9.477,0.8120), t(5.180)=2.142, p=.0831. N=4-5 samples. N.s. not significant (p>.05),*p<.05 C) *daf-8* + Cessation vs. *daf-8* + Water (One-sample t-test: Theoretical mean=1, actual mean=.4398 (95% CI:, -.9426,-.1777), t(3)=4.661, p=.0186. N= 4 samples. *p<.05. D) Relative fold change of *glr-1* mRNA levels in *crh-1* worms relative to wild-type controls (Welch’s t-test: *crh-1* + Water vs. wild-type + Water: Theoretical mean=1, actual mean=2.807 (95% CI:-.7304,4.344), t(3)=2.266,p=.1083; *crh-1* + Cessation vs. Wild-type + Water: Theoretical mean=1,actual mean=1.583, (95% CI:-.5343,1.700), t(3)=1.660,p=.1954). N=4 samples, N.s., not significant (p>.05). E) *crh-1* + Cessation vs. *crh-1* + Water (One-sample t-test: (95% CI:-.8482,.6318), t (4)=.4059, p=.7056)). N=4 samples. N.s., not significant (p>.05). Error bars are SEM.

### CREB-mediated transcription is increased and negatively regulates *glr-1*dependent behaviors after EtOH cessation

Due to our findings that *crh-1* is normally a transcriptional repressor of *glr-1*, and losing *crh-1* is protective against *glr-1* downregulation during EtOH cessation, we next hypothesized that *crh-1* is activated during cessation from chronic EtOH. To address this, we utilized a reporter of *crh-1* transcriptional activity (*pcre::GFP,* Kimura et al., 2002), to track and measure CRH-1/CREB activity in neurons across all treatment conditions. Previous studies demonstrated that this reporter was responsive to starvation (Suo et al., 2006); we repeated these conditions as a positive control for our ability to successfully detect changes in reporter activity (**Fig. 5C**). Untreated negative controls and starved positive controls are validated in **Extended Figure 3-1 A-C**. We found that GFP intensity of both neurons (**Fig. 5A**) and non-neuronal (**Extended data, Fig. 3-1 D, E**) structures after 1h cessation was similar to that of chronic EtOH treatment (**Fig. 5A**). However, the number of GFP+ neurons detected was higher in the cessation group versus the ethanol group (**Fig. 5B**) while number and intensities of non-neuronal structures detected remained the same in both groups (**Extended Data Fig. 3-1E**). Overall, this suggests that neuronal CREB-mediated transcription is increased in 1h cessation.

**Figure 5:**
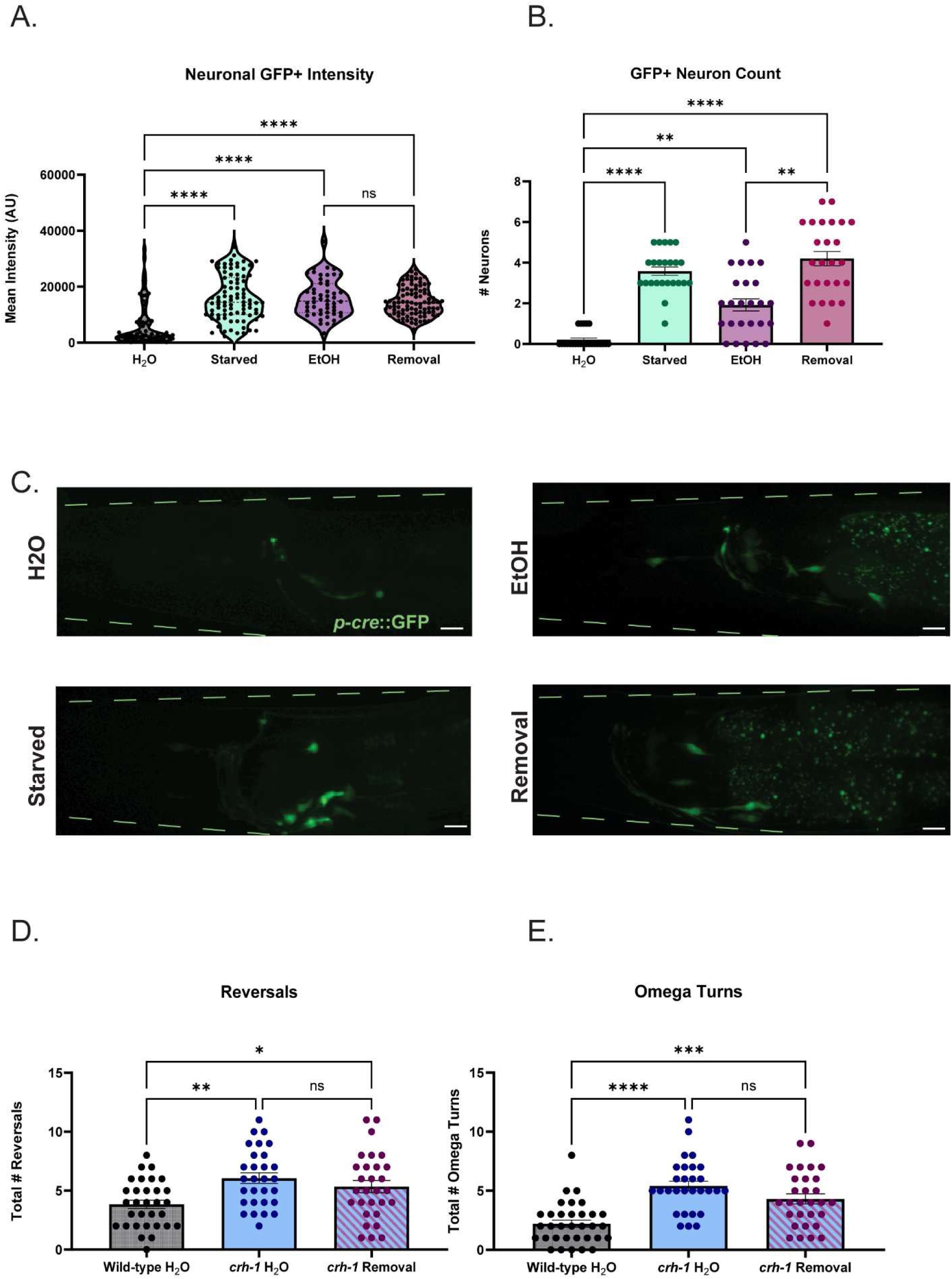
CREB-mediated transcription is increased and negatively regulates *glr-1*-dependent behaviors after EtOH cessation. A) Neuronal GFP intensities of *pcre::GFP* worms across all exposure conditions (Kruskal-Wallis test: p<.0001, 77.69. Dunn’s multiple comparisons test: Mean Rank Water=71.09 vs. Mean Rank Starved=178.8, Z=7.690, p<.0001; Mean Rank Water=71.09 vs. Mean Rank EtOH=180.3,Z=6.855,p<.0001; Mean Rank Water=71.09, Mean Rank vs. Cessation=169.0,Z=7.112,p<.0001; Mean Rank EtOH=180.3 vs. Mean Rank Cessation=169.0, Z=.7482, p>.9999). N=24-25 worms per condition. N.s., not significant (p>.05), ****p<.0001. Error bars are SEM. B) Number of GFP+ neurons detected in *pcre::GFP* worms across all exposure conditions (Kruskal-Wallis test: 60.21, p<.0001. Dunn’s multiple comparisons test:Mean Rank Water=16.35 vs.Mean Rank Starved=66.54, Z=6.205, p<.0001; Mean Rank Water=16.35 vs. Mean Rank EtOH=42.44,Z=3.258, p=.0067; Mean Rank Water=16.35 vs. Mean Rank Cessation=72.02,Z=6.953 ,p<.0001; Mean Rank EtOH=42.44 vs. Mean Rank Cessation=72.02, Z=3.733,p=.0011). N=24-25 worms per condition *p<.05,**p<.007,****p<.0001. Error bars are SEM. C) Representative images of *pcre::GFP* worms across all conditions. 40x objective + 1.5 zoom, scale bars = 10µM. F) Total number of reversals during local search behavior in wild-type, *crh-1*, and *crh-1* worms after water or cessation treatment. (Brown-Forsythe ANOVA; F (2,86)=6.630, p=.0021. Tukey’s multiple comparisons test: Mean Wild-type + Water=3.839 vs. Mean_crh_1_+Water_=6.069 (95% CI: -3.657,-.8037), q(86)=5.030, p=.0018; Mean_Wild-type + Water_=3.839 vs. Mean*_crh-1_*_+Cessation_=5.345 (95% CI: -3.063,.05108), q(86)=3.397, p=.0480; Mean*_crh-1_*_+Water_=6.069 vs.Mean*_crh-1_*_+Cessation_=5.345,(95% CI:-.9727,2.421), q(86)=1.607,p=.4946. N = 29-31 worms per condition. N.s., not significant (p>.05), *p<.05, **p<.005. Error bars are SEM. G) Total number of omega turns during local search behavior in wild-type, *crh-1*, and *crh1* worms after water or 1h cessation (1-way ANOVA, F(2,88)=18.81, p<.0001; Tukey’s multiple comparisons test; Mean_Wild-type + Water_=2.12, vs. Mean*_crh-1_*_+Water_=5.414, (95% CI: 4.474,-1.930), q(88)=8.485, p<.0001; Mean_Wild-type + Water_=2.12 vs. Mean*_crh-1_* _+_ Water=4.310,(95% CI: -3.370,-.8261), q(88)=5.561, p=.0005; Mean*_crh-1_*_+Water_=5.414 vs. Mean*_crh-1_*_+Cessation=_4.310, (95% CI: -.2091,2.416),q(88)=2.834, p=.1172.) N=29-33 worms per condition. N.s., not significant (p>.05), ***p=.0005, ****p<.0001. Error bars are SEM. Extended data:

Because losing *crh-1* is protective against *glr-1* downregulation, we hypothesized that EtOH cessation in *crh-1* mutants would not exhibit behavior deficits that wild-types exhibit in our previously tested *GLR-1* dependent behaviors. However, we found that *crh-1* worms alone perform significantly worse at the ITM time-point compared to wild-type worms in our positive olfactory associative memory assay (**Extended Data Fig. 3-1F**). As a widely expressed transcription factor, CREB plays a significant role in regulation of neuronal function, and this behavioral deficit is likely not solely due to *glr-1*/AMPA receptors. To test the combined roles of CREB and GLR-1 on ITM, we created double mutants of both genes. When we tested the double mutants in our assay, we found losing both *crh-1* and *glr-1* led to significant deficits beginning at 30 minutes after training (short-term associative memory, Extended Data Fig. 3-1G) compared to wild-type worms and single mutants of *crh-1* and *glr-1*. This led us to conclude that the memory deficits observed in *crh-1* mutants are at least partially independent of *glr-1* and would confound our ability to interpret the effects of CRH-1/CREBs role in EtOH cessation mediated memory-deficits. We therefore examined if *crh-1* mutants exhibited resistance to cessation-mediated alterations in local search behaviors, which are a more *glr-1*-specific phenotype. We found that ethanol cessation had no detectable effect on either reversals or omega turns of *crh-1* mutants (**Fig 5D, E**) compared to water-treated controls, indicating that losing *crh-1* is protective against loss of *glr-1* phenotypes. Together these findings suggest that *crh-1* negatively regulates *glr-1* in the context of EtOH cessation and may be linked to deficits in *glr-1*-dependent behaviors in EtOH cessation.

## DISCUSSION

Species have evolved to navigate the world and identify beneficial environments to promote their survival. The behaviors that animals engage in to respond to their environment are plastic and can be modified by experience such as contact with food and odorants, as well as other substances in the wild. Worms can encounter EtOH in the wild potentially through rotting fruit and plant matter (Felix and Duveau, 2012). EtOH is an energy source for worms (Kaptan et al., 2020; Patananan et al., 2015) and low doses extend lifespan in larvae (Yu et al., 2011; Castro et al.,2012). EtOH has molecularly conserved effects across phyla, and can influence behaviors conserved in worms such as state-dependent learning (Lindsay et al., 2022), sensitivity and tolerance (Davies et al., 2003; Davies et al., 2004; Bettinger et al., 2012; Alaimo et al., 2013; Guzman et al., 2022; Mathies et al., 2015; Pandey et al., 2021; Morgan and Sedensky, 1995; Mitchell et al., 2007; Hawkins et al., 2015; Jee et al., 2013; Jee et al., 2024; Johnson et al., 2017), EtOH preference (Lee et al., 2009; Oh et al., 2019) seeking (Salim et al., 2022), and nonassociative memory (Rose et al., 2023; Kokan et al.,2025). In this study, we have leveraged this to model other associative behavioral phenotypes in the context of EtOH cessation.

### ETOH CESSATION, BUT NOT CHRONIC ETOH, DISRUPTS ITM

Here we report that cessation after chronic EtOH exposure causes ITM deficits, whereas continuous chronic EtOH exposure prior to behavioral training has no detectable effect on memory performance in our assay. The resistance to memory deficits in the chronic EtOH group may reflect behavioral tolerance, a form of behavioral plasticity resulting in decreased responses over the course of drug exposure (Kalant, 1998; Pietrzykowsky and Treistman, 2008). Tolerance to EtOH often involves molecular adaptations that function independently of altered pharmacokinetics (i.e. EtOH metabolism). These mechanisms are conserved and include but are not limited to genetic mechanisms (Ghezzi et al., 2004; Davies et al., 2003; Scholz et al., 2000; Scholz et al, 2005; Davies et al., 2004; Thiele et al., 1998), epigenetic modifications (Wang et al., 2007a; Ghezzi et al., 2017), lipid microenvironments (Crowley et al., 2003; Yuan et al., 2008; Bettinger et al., 2012), and alterations in membrane proteins (Liang et al., 2007; Kumar et al., 2003; Davies et al., 2003). Using the robust set of experimental tools available in *C. elegans* to better understand mechanisms of EtOH tolerance in the context of memory deficits will be interesting to study in the future.

Fascinatingly, it was discovered that exposing worms to 400mM EtOH during nonassociative memory training disrupted long-term memory (24h memory) and blocked training-induced changes in *GLR-1* punctal expression (Rose et al., 2023). This contrasts with our experiments, where worms received continuous exposure for 24h prior to memory training and are trained in the absence of EtOH exposure. It is possible that the duration of EtOH exposure during the non-associative memory training in the previous study was insufficient to recruit compensatory mechanisms that would permit normal long-term memory in pre-exposed worms. Together, these studies provide evidence that, not only are active exposure vs. cessation distinct nervous system and behavioral states, but there are also distinctions within the duration of EtOH exposure itself.

EtOH cessation or cessation-specific behavior deficits have been identified in worms including food-based, odor-based, and locomotor assays, where low-dose EtOH rescued some of these behaviors (Mitchell et al., 2010; Scott et al., 2017). In this study, we used a modified version of our protocol and identified a cessation-specific deficit in consolidation of the relationship between an odorant and a food source, increasing the repertoire of cessation-sensitive behaviors in *C. elegans*. Possible reasons for why observed cessation-specific deficits after the worm has adapted to chronic EtOH include that the nervous system may 1) continue to operate at a new baseline as though EtOH is still onboard (it has not readjusted) or 2) is rebounding in response to cessation of the drug. Homeostatic adjustments of neuronal excitability and changes in gene expression are known to occur between chronic EtOH and cessation (Lovinger and Roberto, 2025). As these behaviors are conserved, future investigations in the worm into how the nervous system rapidly responds at a molecular level to cessation will provide a deeper understanding of altered phenotypes after EtOH cessation across species.

### ETOH CESSATION DECREASES *glr-1* EXPRESSION AND FUNCTION

We found that cessation-related ITM deficits may involve *glr-1*. EtOH cessation phenocopies reduced GLR-1 function in both our associative memory assays and in navigation behaviors that are regulated by *glr-1*, and loss of *glr-1* occludes ITM and other behavioral deficits. We discovered that these cessation-dependent behavioral changes correspond with significant downregulation of *glr-1* expression. Overall, our behavior and molecular data point to a decrease in GLR-1/AMPAR expression, and likely a decrease in GLR-1 function, specifically after EtOH cessation, but not during chronic EtOH exposure. It was previously found that cessation from 6h of 200-300mM EtOH led to fewer reversals but a higher frequency of unaccompanied omega-turns during a food-race assay (Mitchell et al., 2010). The increased omega-turns occurred early (5 mins) during the assay. This contrasts with our experimental set-up, where worms are not racing but simply placed on a foodless plate and recorded 7-12 mins following transfer. Factors such as 1) exogenous EtOH exposure, 2) duration of exposure, and 3) behavior-task specific contexts (presence or absence of food) may account for this discrepancy. There are conflicting reports in other models, including mammalian in vitro, in vivo, and human studies as to how AMPARs and glutamate signaling are regulated in cessation. These include instances of increased AMPAR levels and signaling (Christian et al., 2012; Wang et al., 2012; Wang et al., 2015; Gerace et al., 2019; Gerace et al., 2021), no changes in AMPA levels (Williams et al., 2018; Freund and Anderson, 1996; Trevisan et al., 1994; Rudolph et al., 1997; Ferreira et al., 2001), and decreased AMPA expression/function, decreased neuronal excitability, impaired LTP, or glutamate levels (Ulrichsen et al., 1996; Francesconi et al., 2009; Stephens et al., 2005; Pickering et al., 2015; Das et al., 2016; Kryger and Wilce, 2010;Mon et al., 2012; Thoma et al., 2011). Heavy drinking is also associated with lower glutamate levels in the brain (Prisciandaro et al., 2016; Ende et al., 2013; Prisciandaro et al., 2019) although these changes may depend on severity of alcohol-use disorder (Prisciandaro et al., 2020; Ende et al., 2013). These changes in glutamate signaling and neuronal excitability are linked to various behaviors including cessation-related symptoms and seizures (Krupitsky et al., 2007; Brousse et al., 2012; Becker and Mulholland, 2014; Läck et al., 2007), and lower AMPAR levels are linked to lower neuronal excitability, LTP-capability, and memory. There are many possible explanations for these discrepancies – length of cessation, exposure/ EtOH paradigm, brain region, and behavioral investigations, highlighting the difficulty in fully unraveling the interplay between cessation and glutamate receptor signaling leading to behavioral dysfunction.

However, our study is consistent with many of these findings: 1) Chronic EtOH and cessation dysregulates excitatory neurotransmitter signaling, 2) Chronic EtOH and cessation are distinct brain states hallmarked by distinct gene expression patterns, 3) AMPAR expression levels can fluctuate throughout the time-course of cessation. One hypothesis for this observed “undershoot” of GLR-1 expression and function in EtOH cessation in our study may reflect homeostatic re-balancing of excitation/inhibition ratios. Indeed, there is evidence that compensatory mechanisms regulate neuronal excitability in chronic EtOH and cessation-including those involving conserved synaptic proteins and ion channels (Mulholland et al., 2011; Welsh et al., 2011; Ghezzi and Atkinson, 2011; Scott et al., 2017; Lovinger and Roberto, 2025; Taylor et al., 2024). It will be interesting to determine if GLR-1 expression changes lead to alterations in synaptic GLR-1, or if other mechanisms of GLR-1 regulation are also disrupted by EtOH cessation. We can conclude that GLR-1 depletion we observe following EtOH cessation likely yields insufficient amounts to allow for ITM consolidation and normal navigation behaviors during local search behavior.

### THE ROLE OF CRH-1/CREB IN GLR-1/AMPAR REGULATION IN CESSATION

We found *crh-1*/CREB prevents both *glr-1* downregulation and some *glr-1* dependent behavior deficits in cessation, and a cessation-dependent increase in numbers of neurons exhibiting CRH-1/CREB reporter activity. These results suggest CREB-mediated transcription is increased in EtOH cessation and may regulate cessation-dependent changes in *glr-1* expression and function. Normally, CREB is required for long-term memory, and is not required for short-term memory across species (Lamprecht, Hazvi, and Dudai, 1997; Frank and Greenberg, 1994; Carew, 1996); Lakhina et al., 2015; Kauffman et al., 2010; Arey et al., 2018). In long-term memory, CREB’s activation and interaction with co-activators leads to persistent changes affecting gene expression, and synapse dynamics following neuronal activity and associative learning (Nonaka et al., 2014; Deisseroth et al., 1996; Hirano et al., 2016; Parra-Damas et al., 2017; Kandel, 2012). We hypothesize that EtOH cessation-related CREB activity in more neurons could result in aberrant upregulation of plasticity-related genes, which could alter synaptic connectivity leading to disrupted behaviors. Interestingly, previous studies in rats, mice, and flies found increases in CREB or phosphorylated CREB in cessation from EtOH (Pandey et al., 2001; Bison and Crews 2003; Kurokawa et al.,2013; Shibasaki et al., 2011; Qiao et al., 2018; Wang et al., 2007a; Freeman et al., 2013), and other drugs of abuse (Fijal et al., 2015; Navarro-Zaragoza et al., 2017). CREB is linked to many behaviors in cessation including anxiety, EtOH consumption, preference, spatial memory, and tolerance (Pandey et al.,1999; Pandey et al., 2004; Qiao et al., 2018; Dominguez et al., 2016; Kurokawa et al., 2013; Wang et al., 2007a). The complexity of the relationship between CREB and EtOH may be due to the same discrepancies described above for AMPARs. Overall, our findings agree with the literature that phases of EtOH exposure and cessation recruit CREB-dependent transcription, with additional findings that loss of CREB protects against some *glr-1*-dependent behavior deficits in EtOH cessation. Interestingly, the interaction between CRH-1/CREB and *glr-1* is likely not due to direct transcriptional regulation. There is not a known CRH-1 binding site on *glr-1*, so why might CREB be activated during EtOH cessation to reduce *glr-1* expression? Given the link between enhanced glutamate signaling in cessation in other models, it is possible CREB activation protects against the threat of excitotoxicity by tamping down *glr-1* expression by recruiting neuroprotective transcripts. In models of ischemia, CREB activation protected against cell-death, as did blocking the calcium-permeability of AMPARs (Liu et al., 2004). In worms, CREB and one of its co-activators, CRTC-1, protect against glutamate toxicity and neuron death (Feldmann et al., 2019). In addition, CREB activity is increased in models of oxidative stress by way of MAP kinases (Németh et al., 2025; Zhi et al., 2020; Yin et al., 2022), which are regulated by EtOH treatment (Németh et al., 2025; Agoglia et al., 2015; Ku et al., 2007) and regulate memory (Impey, Obrietan, and Storm, 1999; Morel et al., 2018; Zhao et al., 2022). In addition, GLR-1 trafficking is impacted by reactive oxygen species (Doser, Knight, and Deihl, 2020). Therefore, memory ability may be temporarily reduced to maintain long-term neuronal health. Future experiments should investigate synaptic GLR-1 dynamics in wild-type and *crh-1* backgrounds in the context of EtOH cessation.

Overall, this study provides evidence that EtOH cessation disrupts post-synaptic glutamatergic signaling mechanisms, potentially by way of CREB-dependent transcriptional regulation, leading to deficits in the consolidation of positive associative olfactory memories. *C. elegans* will be useful to further unravel the relationship between environmental exposure and EtOH exposure states, memory, and behavioral phenotypes.

## Acknowledgements

We thank the Stavoe lab, Eddie Pietryk III, and Catherine Stuart for helpful discussions on microscopy experiments and immunoblotting techniques, and Ashley Hayden for early experimental assistance. We thank all Arey lab members for their detailed feedback on the manuscript.

## Figure Legends

**Extended data Figure 1-1:**
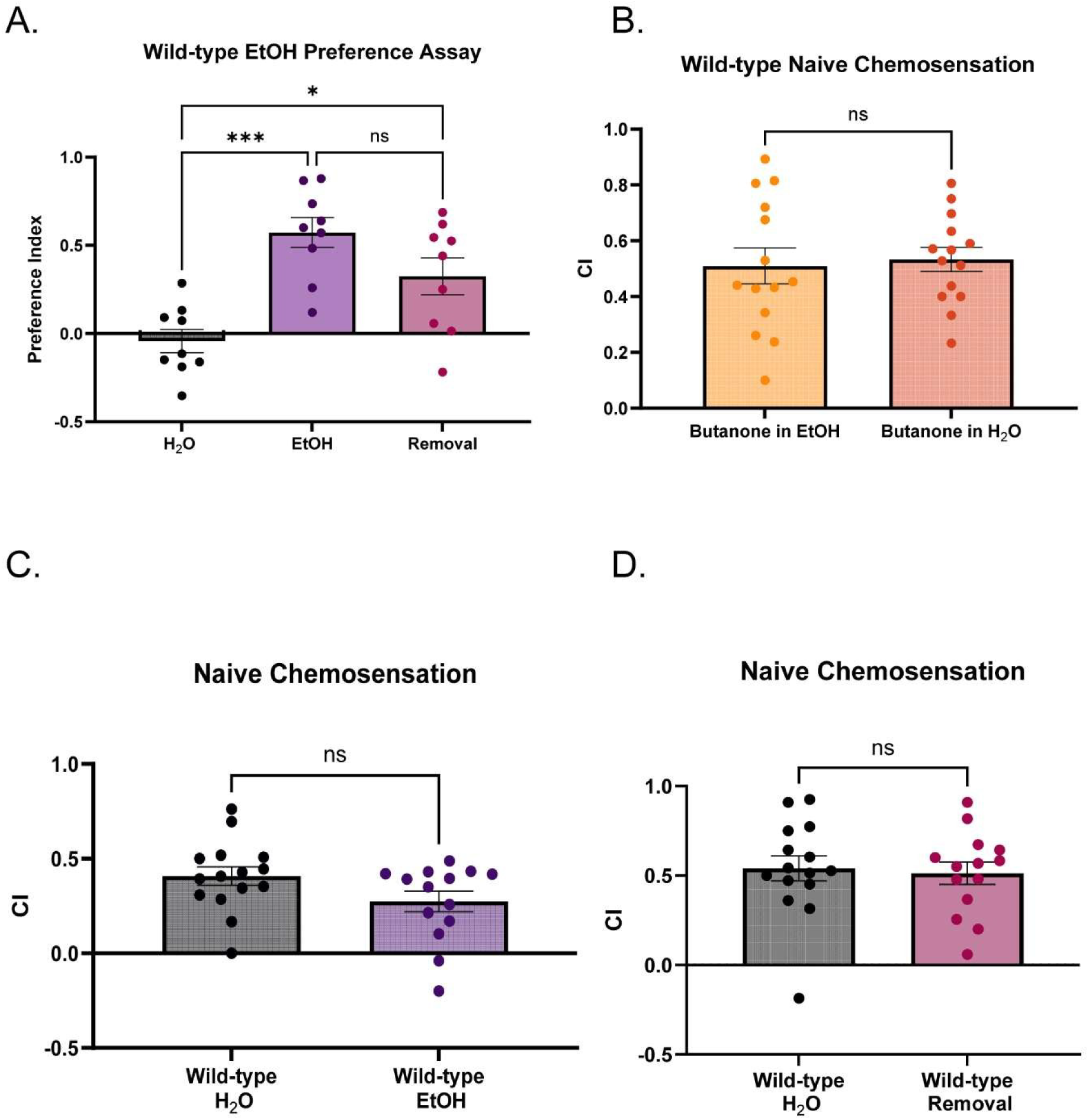
A) EtOH preference indices across treatment conditions in wild-type worms. (Brown-Forsythe ANOVA 12.70(2,21.29), p=.0002. Dunnett’s T3 multiple comparisons test: Mean_Water_=-.04247 vs. Mean_EtOH_=.5729,(95% CI: -.9036,-.3273), t(15.10)=5.699, p=.0001;Mean_Water_=-.04247 vs.Mean_Cessation_=.3245,(95% CI: -.7008,-.03327), t(13.53)=2.959, p=.0298; Mean_EtOH_=.5729 vs. Mean_Cessation_=.3245, (95% CI: .1120,.6087),t(15.36)=1.839, p=.2266). N=9 plates per condition. N.s., not significant (p>.05), *p<.05, ***p<.005. Error bars are SEM. B) Naïve chemotaxis indices of wild-type worms with the test odorant (10% butanone) diluted in EtOH vs. water. (Welch’s t test: Mean_Butanone+Water_=.5328 vs. Mean_Butanone+EtOH_=.5097 (95% CI: -.1371,.1834), t(22.75)=.2990, p=.7677). N=14 plates per condition. N.s, not significant(p>.05). Error bars are SEM. C) Naïve chemotaxis indices of water-treated and EtOH-treated wild-types (Welch’s t test: Mean_Water_=.4077 vs. Mean_EtOH_=.2739 (95% CI: -.2839,.01611), t(26.41) =1.833, p=.078). N=14-15 plates per condition, N.s., not significant (p>.05). Error bars are SEM. D) Naïve chemotaxis indices of water-treated and cessation-treated wild-types (Welch’s t test: Mean_Water_ =.5403, Mean_Cessation_ =.5132(95% CI: -.2189,.1646), t(26.84)=.2907, p=.7735). N=14-15 plates per condition, N.s., not significant (p>.05). Error bars are SEM.

**Extended data Figure 2-1:**
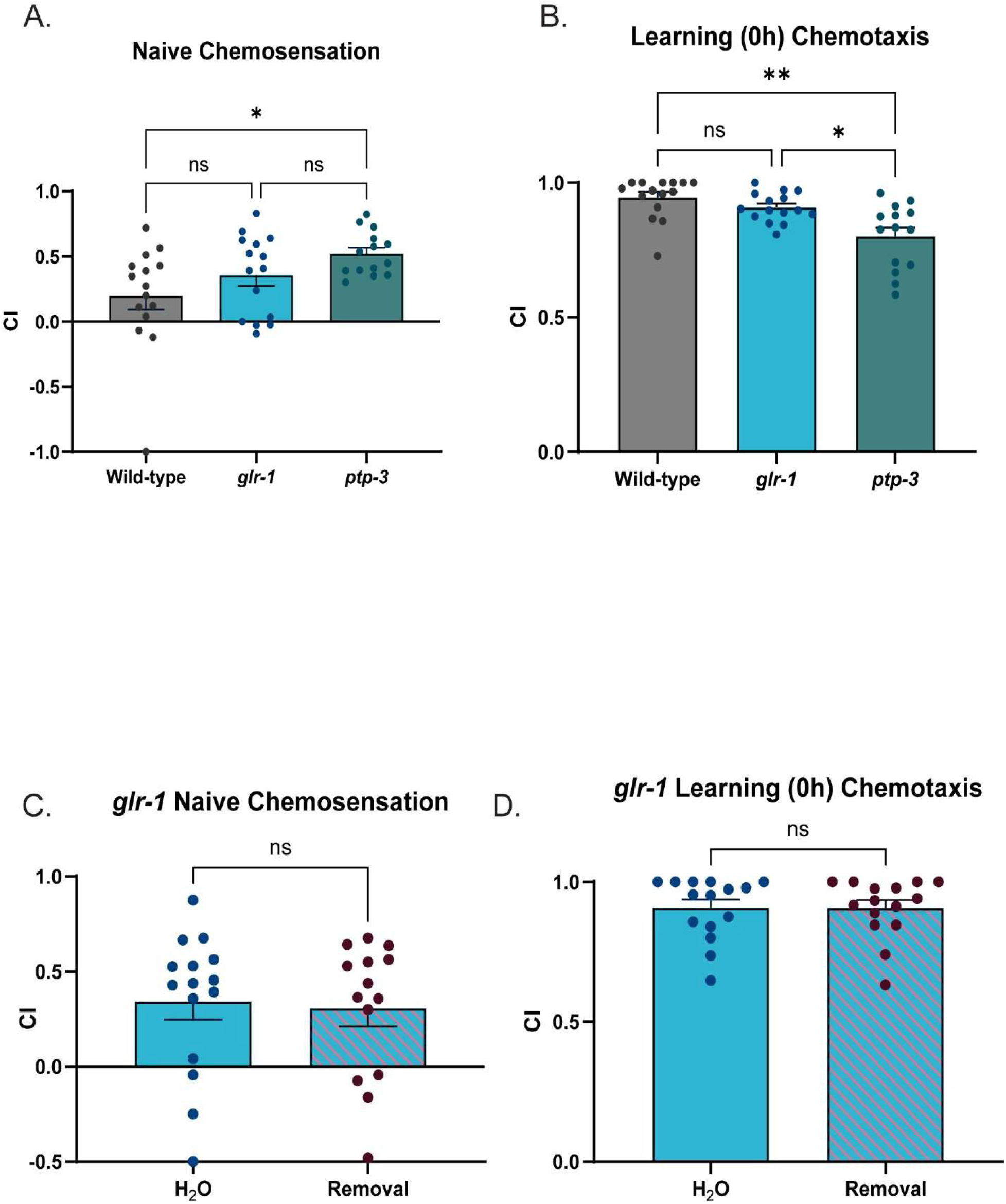
A) Naïve chemotaxis indices of wild-type, *glr-1*, and *ptp-3* worms. (Brown-Forsythe ANOVA: F=4.019(2,31.54), p=.0279. Dunnet’s T3 Multiple comparisons test: Mean_Wild_type=.1960 vs.Mean*_glr-1_*=.3548 (95% CI:-.4946,.1771) t(26.13)=1.202, p=.5515; Mean_Wild_type=.1960 vs. Mean*_ptp-3_*=.5222 (95% CI: -.6242,.02189),t(18.94)=2.852, p=.0296; Mean*glr-1*=.3548 vs. Mean*_ptp-3_*=.5222 (95% CI:-.4036,.06872), t(21.94)=1.825, p=.2194). N=14-15 plates per condition. N.s., not significant (p>.05), *p<.05). Error bars are SEM. B) Learning (0h) chemotaxis indices of wild-type, *glr-1*, and *ptp-3* worms. (BrownForsythe ANOVA, F=10.16(2,27.5), p=.0005. Dunnett’s T3 multiple comparison’s test: Wild-type vs. *glr-1*: Mean_Wild-type_= .9457, Mean*_glr-1_*=.9082(95% CI: .02454,.09953),t(25.11)=1.542,p=.3465; Wild-type vs. *ptp-3*: Mean_Wild-type_=.9457, Mean*_ptp-3_*=.8006, (95% CI:.04662,.2437), t(21.66)=3.792,p=.0030; *glr-1* vs. *ptp-3*: Mean_glr_1=.9082, Mean*_ptp-3_*=.8006 (95% CI:.01454,.2008),t(17.64)=3.028,p=.0211). N=14-15 plates per condition. N.s., not significant (p>.05), *p<.05, **p<.005. Error bars are SEM. C) Naïve chemotaxis indices of *glr-1* worms treated with water or EtOH cessation (Welsh’s t test: Mean_Water_:.3436, Mean_Cessation_:.3068 (95% CI: -.3152,.2416), t (26.98)=.2713, p=.7882). N=14-15 plates per condition. N.s., not significant (p>.05). Error bars are SEM. D) Learning (0h) chemotaxis indices of *glr-1* worms treated with water or EtOH cessation (Welsh’s t test: Mean_Water_=:.9077, Mean_Cessation_=.9075 (95% CI: -.08156,.08107), t(27.94) =.006104, p=.9952). N=14-15 plates per condition. N.s., not significant (p>.05). Error bars are SEM.

**Extended data Figure 3-1:**
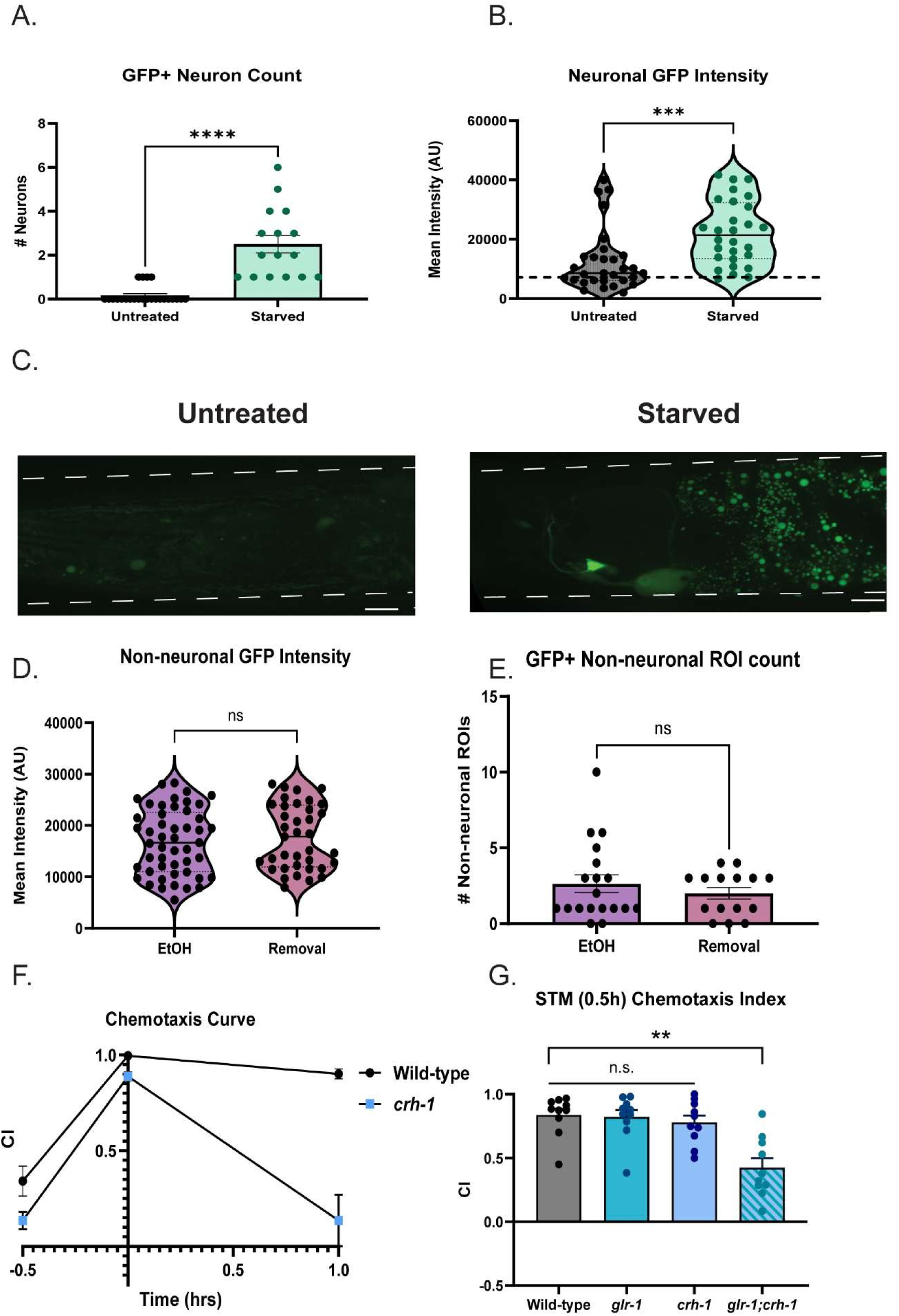
A) Number of GFP+ ROIs detected in *pcre::GFP* negative and positive controls (4h starved). (Mann-Whitney test, U=12 M_Untreated_=0.00, M_Starved_=2.000. N=16-24 worms per condition. ****p<.0001. Error bars are SEM. B) Neuronal GFP intensities of *pcre::GFP* untreated and 4h starved controls. (MannWhitney test, U=181 M_Untreated_ = 8567, M_Starved_ = 21364, p<.0001). N=28-30 worms per condition. ***p=.0001 Error bars are SEM. C) Representative images of *pcre::GFP* untreated vs. 4h starved controls. 100X objective. Scale bars = 10µM. D) Number of non-neuronal GFP+ ROIs detected in EtOH-treated or cessation treated *pcre:;GFP* worms (Mann-Whitney test: U=131, M_EtOH_=1.00, M_Cessation_=3.00, p=.6934). N=15-20 worms per condition. N.s., not significant (p>.05). Error bars are SEM. E) Non-neuronal GFP intensities in EtOH and 1h cessation treated *pcre::GFP* worms (Mann-Whitney test: U=680, M_EtOH_=18760, M_Cessation_=18658, p=.7626). N=15-20 worms per condition. Error bars are SEM. F) Chemotaxis curve of wild-type vs. *crh-1*, double mutants across olfactory memory assay (Mixed-effects analysis: Time x Genotype, p<.0001. Bonferroni’s multiple comparisons test: Naïve: Mean_Wild-type_=.3413, Mean*_crh-1_*=.1352 (95% CI: -.03962,.4518), t(14.27)=2.274, p=.1168; 0h: Mean_Wild-type_=.9965, Mean*_crh-1_*= .8883 (95% CI: .04515,.1713) t (9.228) =5.006, p=.0020;1h: Mean_Wild-type_=.9016, Mean*_crh-1_*=.1354 (95% CI: .3623,1.170), t(8.608)=5.621, p=.0011. N=9-10 plates per condition. N.s., not significant (p>.05), **p<.005. Error bars are SEM. G) STM (0.5h) chemotaxis of wild-type vs. *crh-1, glr-1* single mutants and *glr-1;crh-1* double mutants. Brown-Forsythe ANOVA: F=11.10(3,32.35),p<.0001. Dunnett’s T3 multiple comparisons: Mean_Wild-type_=.8377 vs. Mean*_glr-1_*=.8228 (95% CI:-.2021,.2319), t(17.82)=.2006,p>.9999; Mean_Wild-type_=.8377 vs. Mean*_crh-1_*=.7796 (95% CI: .1558,.2719),t(17.90)=.7952,p=.9590; Mean_Wild-type_=.8377 vs. Mean*_glr-1_*_;*crh-1*_= .4249 (95% CI: .1484,.6772),t(15.75)=4.636,p=.0016; Mean*_glr-1_*=.8228 vs. Mean*_crh-1_*= .7796 (95% CI:.1816,.2680),t(17.99)=.5629,p=.9926; Mean*_glr-1_*=.8228 vs. Mean*_glr-1_*_;*crh-1*_=.4249 (95% CI:.1264,.6695),t(16.63)=4.320,p=.0027; Mean*_crh-1_*=.7796 vs. Mean*_glr-1_*_;*crh-1*_=.4249 (95% CI:.08376,.6257),t(16.40)=3.887,p=.0076 N=10 plates per condition. N.s., not significant, **p<.008. Error bars are SEM.

